# Salmonella enterica serovar Typhimurium SPI-1 and SPI-2 shape the transcriptional landscape of epithelial cells in a human intestinal organoid model system

**DOI:** 10.1101/2020.10.05.327551

**Authors:** Anna-Lisa E. Lawrence, Basel H. Abuaita, Ryan P. Berger, David R. Hill, Sha Huang, Veda K. Yadagiri, Brooke Bons, Courtney Fields, Christiane E. Wobus, Jason R. Spence, Vincent B. Young, Mary X. O’Riordan

## Abstract

The intestinal epithelium is a primary interface for engagement of the host response by foodborne pathogens, like *Salmonella enterica* serovar Typhimurium (STm). While interaction of STm with the mammalian host has been well studied *in vitro* in transformed epithelial cell lines or in the complex intestinal environment *in vivo*, few tractable models recapitulate key features of the intestinal epithelium. Human intestinal organoids (HIOs) contain a polarized epithelium with functionally differentiated cell subtypes, including enterocytes and goblet cells. HIOs contain luminal space that supports bacterial replication and are more amenable to experimental manipulation than animals while more reflective of physiological epithelial responses. Here we use the HIO model to define transcriptional responses of the host epithelium to STm infection, also determining host pathways dependent on *Salmonella* Pathogenicity Island-1 (SPI-1) and -2 (SPI-2) encoded Type 3 secretion systems (T3SS). Consistent with prior findings, we find that STm strongly stimulates pro-inflammatory gene expression. Infection-induced cytokine gene expression was rapid, transient and largely independent of SPI-1 T3SS-mediated invasion, likely due to continued luminal stimulation. Notably, STm infection led to significant down-regulation of host genes associated with cell cycle and DNA repair, an effect that required SPI-1 and SPI-2 T3SS. The transcriptional profile of cell cycle-associated target genes implicates multiple miRNAs as likely mediators of STm-dependent cell cycle suppression. These findings from Salmonella-infected HIOs delineate common and distinct contributions of SPI-1 and SPI-2 T3SSs in inducing early host responses during enteric infection and reveal host cell cycle as a potential target during STm intracellular infection.

**Importance:** *Salmonella enterica* serovar Typhimurium (STm) causes a significant health burden worldwide, yet host responses to initial stages of intestinal infection remain poorly understood. Due to differences in infection outcome between mice and humans, evaluating physiological host responses driven by major virulence determinants of *Salmonella* have been difficult to date. Here we use the 3D human intestinal organoid model to define early responses to infection with wildtype STm and mutants defective in the SPI-1 or SPI-2 Type 3 secretion systems. Both secretion system mutants show defects in a mouse model of oral *Salmonella* infection but the specific contributions of each secretion system are less well understood. We show that STm upregulates pro-inflammatory pathways independently of either secretion system while downregulation of host cell cycle pathways is dependent on both SPI-1 and SPI-2. These findings lay the groundwork for future studies investigating how SPI-1- and SPI-2-driven host responses affect infection outcome and show the potential of this model to study host-pathogen interactions with other serovars to understand how initial interactions with the intestinal epithelium may affect pathogenesis.

## Introduction

Enteric bacterial infections constitute a major human disease burden worldwide, with *Salmonella* species accounting for the most hospitalizations in outbreaks with a confirmed cause. In total, *Salmonella* causes an estimated 1.35 million infections in the US alone (1). Enteric infections occur in a complex and highly dynamic environment that traverses the distinct landscapes of the gastrointestinal tract. Relevant to understanding infection are the host processes that shape physicochemical properties of the intestine, including regulation of pH and nutrient absorption, the epithelial layer, which establishes a barrier using epithelial tight junctions, mucus and antimicrobial peptides, the microbiome and the pathogen itself. While animal models are valuable *in vivo* approaches to understand enteric infections, these models suffer from two major limitations. First, the complexity of the mammalian intestine makes finely controlled experimental manipulation and observation challenging. Second, the physiology of the intestine in different organisms can differ sharply, i.e., mice rarely exhibit diarrhea upon infection by pathogens that would cause diarrhea in humans.

*Salmonella enterica* serovar Typhimurium (STm) infection is a prime example of this disease difference between humans and mice. While STm infection is most commonly associated with self-limiting gastroenteritis in otherwise healthy humans, it causes systemic acute disease in C57Bl/6 mice (naturally Nramp-deficient) or chronic disease in Nramp-sufficient mouse strains (2, 3). To interrogate molecular mechanisms of host:pathogen interactions during intracellular STm infection, many previous studies have relied on transformed human cells, such as the HeLa cervical epithelial cell line or Caco-2 intestinal epithelial cell line, or primary mouse cells like embryonic fibroblasts or macrophages. These cell culture systems have revealed much about STm infection, yet do not recapitulate several key features likely to be important during STm enteric infection. These include the continued presence of STm in the lumen, known to be an environment that supports robust replication, and interaction with non-transformed intestinal epithelial cells (IEC) which have specific properties, like mucus secretion or controlled cell cycle regulation. Thus, elucidating the cellular and molecular basis of STm:epithelial interactions in non-transformed human epithelial cells will improve our understanding of aspects of infection that may be relevant to human disease.

In the last decade, human intestinal organoid (HIO) systems have been developed to enable study of more complex IEC characteristics. These organoids are differentiated from non-transformed human pluripotent stem cell lines such as embryonic or induced pluripotent stem cells (ESC/iPSC), and form 3D cyst-like structures delineated by polarized epithelium with a mesenchymal layer surrounding a luminal space (4). HIOs contain multiple epithelial cell subsets, including enterocytes and goblet cells (4). A previous study characterized the global transcriptional profile of WT STm-infected HIOs using human induced pluripotent stem cells (hiPSC), and demonstrated that this IEC model could support STm infection (5). Their results established that the IEC transcriptional response to WT STm infection from the apical or basolateral route was dominated by pro-inflammatory innate immune signaling pathways. Further studies have demonstrated that HIOs can support survival and or replication of both pathogenic and commensal bacteria, and that commensal organisms, like *Escherichia coli* (ECOR2), stimulate epithelial maturation and barrier function (6, 7).

Here we use HIOs derived from the H9 human embryonic stem cell line to define the host transcriptional response to infection by the commonly used laboratory strain STm SL1344 compared to isogenic mutants lacking functional SPI-1 or SPI-2 Type 3 secretion systems (T3SS); major virulence determinants of STm, which secrete effector proteins into the host to mediate cellular invasion and remodeling of host processes (8, 9). We find that STm-infected HIOs recapitulate some key aspects of intracellular infection as reported, but additionally that the continued presence of luminal bacteria drives a robust epithelial innate immune response even when STm invasion is minimal. Moreover, our results show that WT STm infection reduces transcript levels of genes involved in cell cycle regulation and DNA repair. These findings underscore the value of the HIO model for studying STm by validating characteristic host responses to infection by WT or mutant STm, as well as revealing new infection-induced host pathways.

## Results

### Luminal *Salmonella* Typhimurium replicate within HIOs and invade HIO epithelial cells

To better recapitulate the *in vivo* human intestinal epithelial response to *Salmonella* infection, we used the 3-dimensional HIO model that allows longer-term bacterial-host interactions compared to traditional cell lines by maintaining the bacteria in the luminal space throughout the course of infection. STm was inoculated into the HIO lumen by microinjecting each HIO with ∼10^3^ colony forming units (CFU), or PBS as a control **(Fig. 1A)**. HIOs were allowed to recover for 2h prior to 15 min treatment with medium containing 100 μg/mL gentamicin to kill bacteria that were introduced into the culture medium during microinjection. Subsequently, HIOs were cultured in medium containing 10 μg/mL gentamicin for the remainder of the infection to prevent replication of STm outside the HIOs. To confirm that STm replication could take place within HIOs, HIOs were injected with STm harboring the pGEN plasmid encoding the fluorescent protein DsRed (10), and bacterial burden was monitored by live fluorescence microscopy **(Fig 1B,C)**. Fluorescence intensity substantially increased by 24h post-infection (pi) indicating that STm replicated within the HIOs and replication appeared to occur predominantly in the lumen. Histological analysis of HIO sections revealed that luminal STm invaded intestinal epithelial cells and migrated to the basolateral side **(Fig. 1D)**. Notably, invasion did not occur uniformly across the HIO as not all cells became infected. Additionally, infection did not appear to cause major structural damage to the HIO as viewed by H&E staining, but infection was accompanied by increased mucus production on the luminal surface of the epithelial barrier **(Fig. S1)**. To further quantify bacterial burden, we enumerated total bacterial CFU per HIO and found a 3-log increase from the 2.5h to 24h time points **(Fig. 1E)**. Consistent with previous reports, invasion was largely dependent on the STm type III secretion system (T3SS) encoded on pathogenicity island 1 (SPI-1) (11), as an inframe deletion in a structural gene of the T3SS apparatus (Δ*orgA*) drastically reduced intracellular CFU **(Fig. 1F)**. Together these results demonstrate that the HIO model supports robust luminal and intracellular replication of STm, and that invasion of HIO epithelial cells is dependent on T3SS-1.

**Figure 1:**
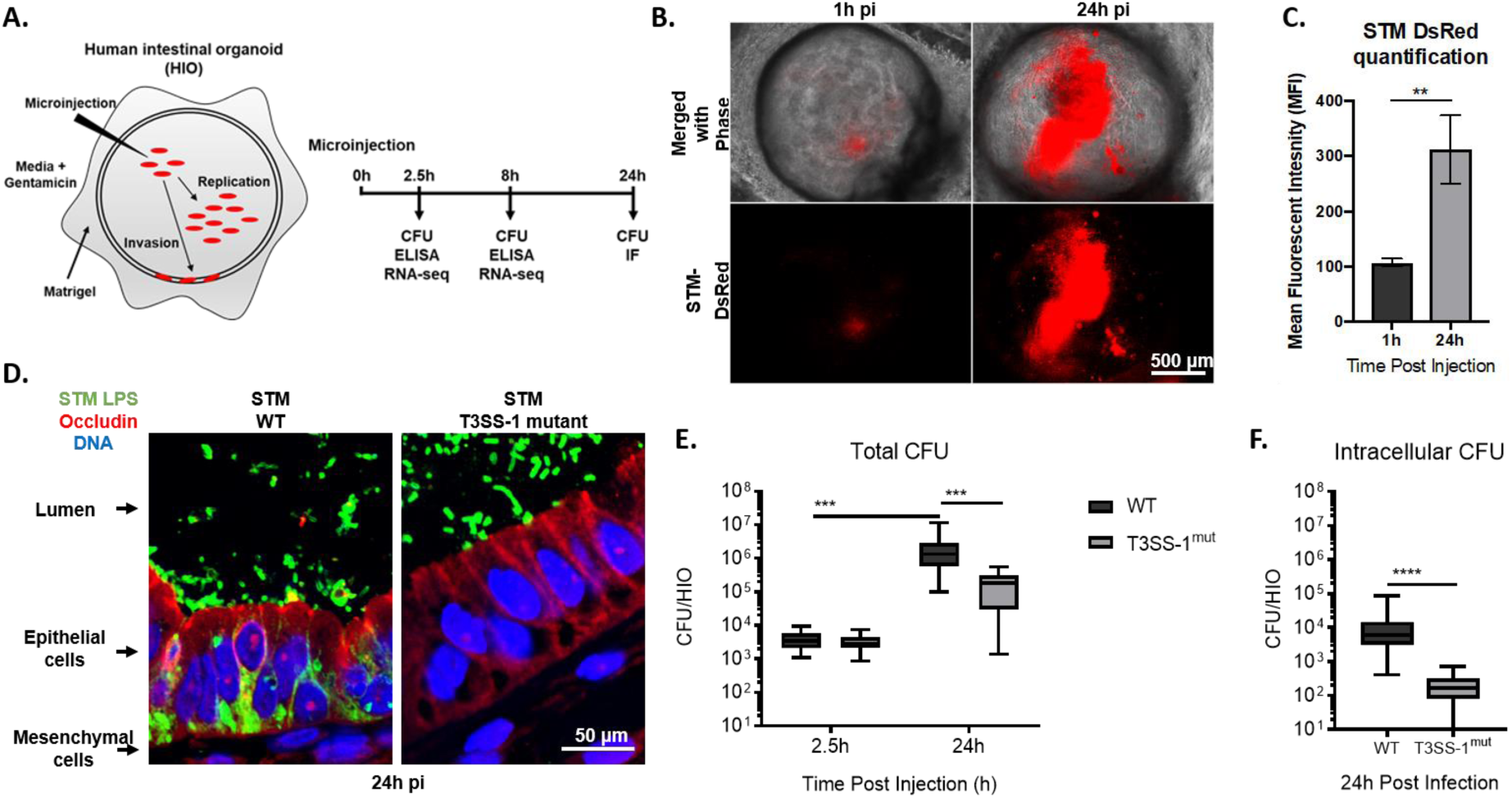
WT STm replicates within the lumen of HIOs and invades IECs dependent on T3SS-1. (A) Diagram of experimental protocol. (B) Fluorescence microscopy of HIOs injected with STm-DsRed, a strain that harbors the pGEN plasmid encoding Red fluorescence protein (DsRed) (10). (C) Quantification of Fig.1B. n = 3 biological replicates. Error bars represent SD. p = 0.0047 by unpaired t test. (D) Immunofluorescence of HIO sections infected with STm WT (left) and STm T3SS-1^mut^ (right). (E) Total bacteria in HIOs at 2.5 and 24h post injection. n=16 biological replicates. Whiskers represent min and max values. Significance calculated by two-way ANOVA. (F) Intracellular bacteria in HIOs at 24h post injection. n > 31 biological replicates. Whiskers represent min and max values. Significance calculated by Mann-Whitney test.

### Kinetic analysis of HIO transcriptional profiles define the acute response to *Salmonella* infection

To gain insight into global HIO transcriptional responses stimulated by STm infection and to define the relative contributions of the major virulence determinants, T3SS-1 and -2, we performed RNA sequencing (RNA-seq) at 2.5h and 8h pi with HIOs microinjected wit h PBS, WT STm or isogenic mutants in Δ*orgA* (T3SS-1^mut^) and Δ*ssaV* (T3SS-2^mut^). Microinjection with each of these strains yielded similar levels of total bacteria at 2.5h while the T3SS-1^mut^ strain showed significantly reduced levels of total and intracellular bacteria at 8h **(Fig. S2)**, consistent with a role for the T3SS-1 in invasion. Principal component analysis (PCA) showed that all infected HIOs displayed markedly different transcriptional profiles than those injected with PBS **(Fig. 2A)**. Notably, sample clustering occurred primarily by time post infection because 2.5h and 8h infected HIOs segregated from each other along the first principal component (x-axis). This difference accounted for 40% of the total variance and suggested that time post infection is a greater determinant of transcriptional variance than mutations in the pathogen. Similar patterns were observed by Pearson’s correlation clustering, which showed clustering of 2.5h and 8h samples **(Fig. 2B)**. In addition, the Pearson’s correlation heat map showed that HIOs infected with the invasion-defective T3SS-1^mut^ segregated away from samples infected with WT STm and the T3SS-2^mut^ at 2.5h pi while at 8h pi, HIOs infected with WT STm separated from both mutants. These data suggest that the T3SS-2^mut^ is attenuated later in infection compared to wild type; a time point at which bacteria have invaded the epithelium, and the T3SS-2 is thought to be active to maintain intracellular infection (8, 12, 13). Using differential expression analysis, we found that HIOs injected with any of the 3 strains of STm resulted in similar numbers of significant gene changes (p-value < 0.05) at 2.5h pi compared to PBS controls, suggesting that the early HIO response is driven primarily by luminal bacteria **(Fig. 2C, Table S1, S2)**. In contrast, at 8h both T3SS mutant strains induced fewer significant gene changes than WT suggesting that T3SS-1 and -2 effectors may be required for STm-induced responses later during infection **(Fig. 2D)**.

**Figure 2:**
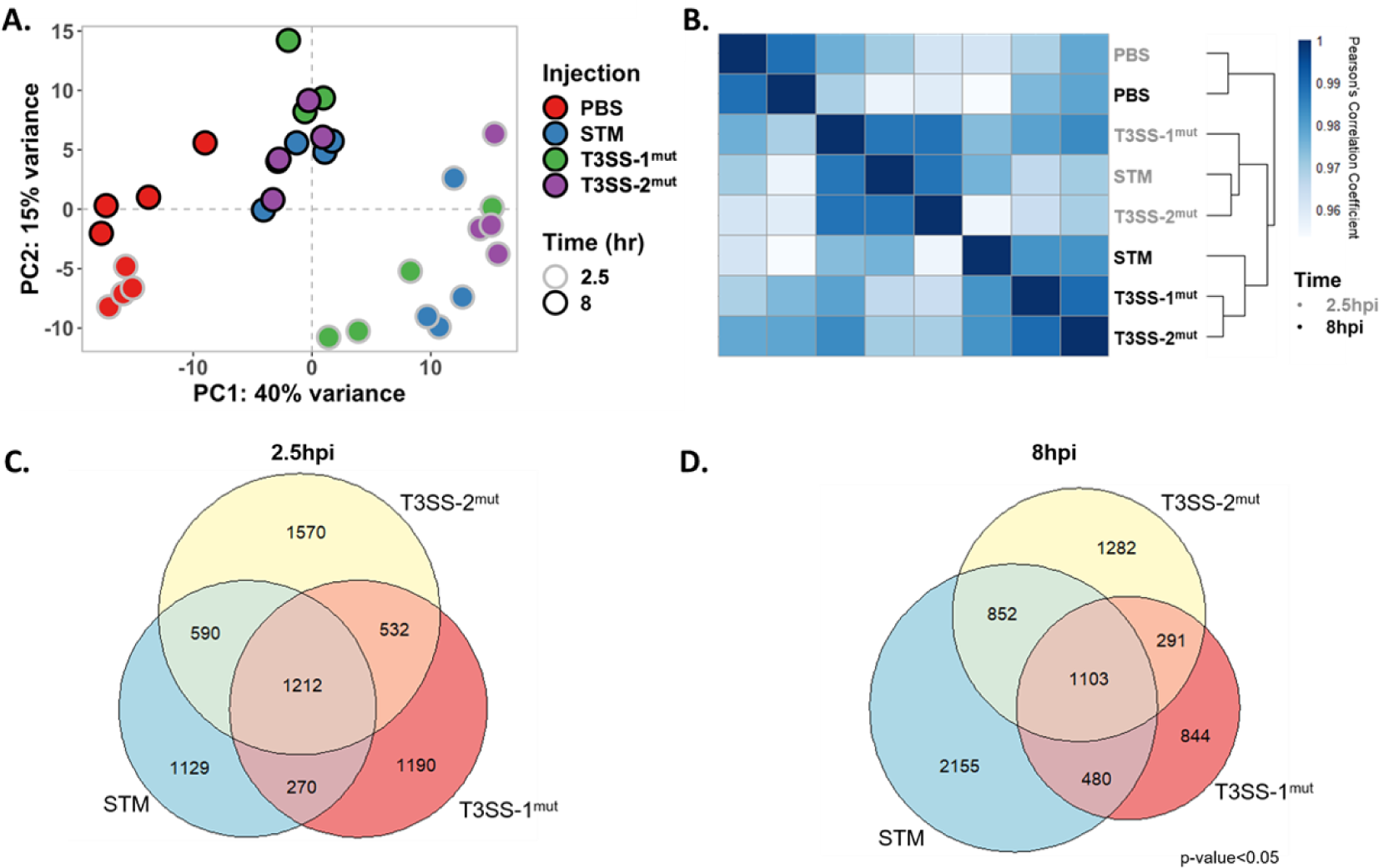
HIOs mount an acute transcriptional response to *Salmonella* infection. (A) Principal component analysis of HIOs injected with STm T3SS mutants. Each circle represents a biological replicate. (B) Sample distance plot of each HIO condition at 2.5h (gray) and 8h (black) post injection. Sample distance calculated from normalized gene counts across 4 biological replicates. (C-D) Euler diagram comparison of gene changes in each HIO condition relative to PBS injected HIOs at (C) 2.5h and (D) 8h post injection. Genes were filtered by p-value < 0.05.

### Immune pathways and cell cycle pathways are inversely regulated during *Salmonella* infection

To determine which pathways drive the epithelial response to STm infection, we performed pathway enrichment analysis from the Reactome database **(Table S3A,B and S4A,B)**. Clustering of sub-pathways into major cellular processes in the Reactome database indicated that the majority of up-regulated pathways in all three infection conditions clustered into immune response and signal transduction processes **(Fig. 3A)**. We examined individual pathway enrichment by gene ratio (fraction of genes in a pathway that were significantly changed) and the -log_10_(p-value) to identify pathways modulated by STm infection and dependence on T3SS-1 or T3SS-2. To our surprise, we observed similar gene ratios between infection with WT STm and the two T3SS mutants in several cytokine signaling pathways, including genes encoding IL4, IL17 and IL10 **(Fig. 3B top)**. These results are in contrast to previous reports that T3SS-1-dependent invasion strongly contributes to the inflammatory response including upregulation of cytokines such as *IL8* (14–16). However, distinct from most tissue culture infection models, the HIO model system features sustained epithelial interaction with both luminal and intracellular *Salmonella*, pointing to a strong contribution of luminal bacteria in triggering early inflammation. Importantly, not all inflammatory pathways were equally enriched in all 3 infection conditions; innate immune signaling pathways, including Toll-like Receptor (TLR) signaling cascades were less enriched in T3SS-1^mut^-infected HIOs at 2.5h pi, and in both T3SS-1^mut^- and T3SS-2^mut^-infected HIOs compared to WT at 8h pi suggesting that modulation of these pathways is enhanced by intracellular infection **(Fig. 3B middle)**.

**Figure 3:**
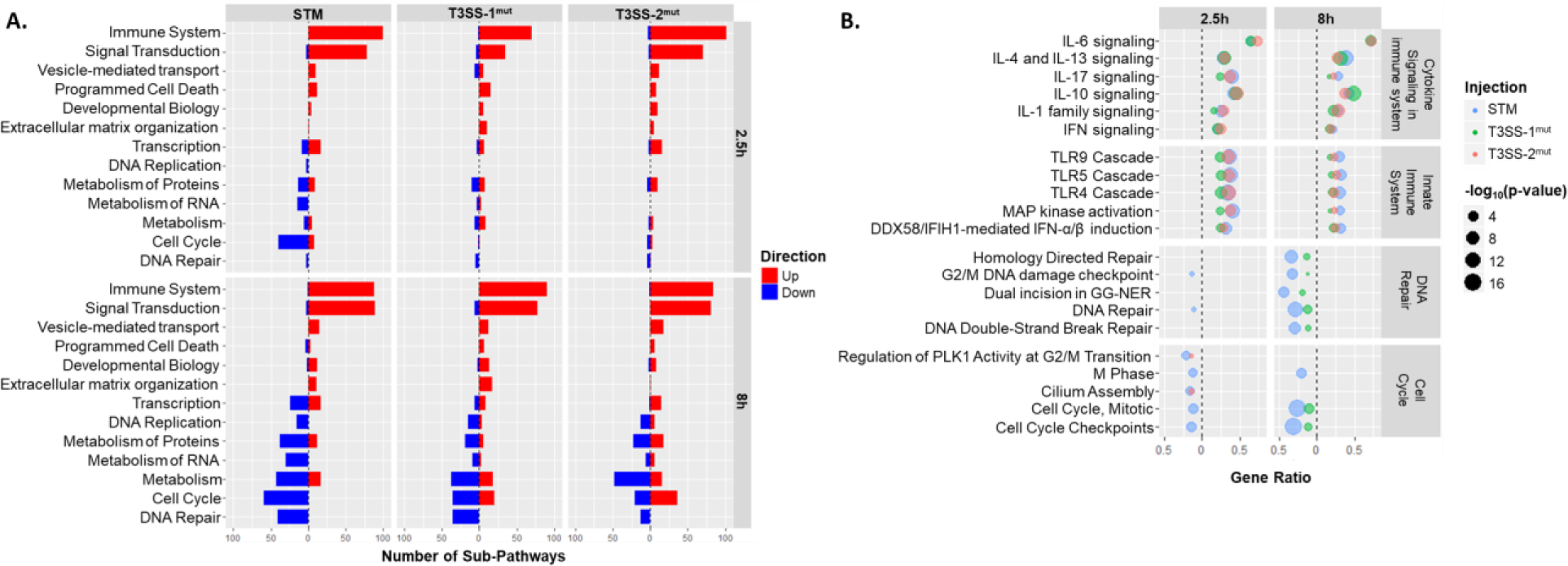
Reactome pathway enrichment reveals upregulation of immune system pathways and downregulation of cell cycle and DNA repair pathways. (A) Number of sub-pathways clustering into major Reactome pathways. Significantly up-regulated (red) or down-regulated (blue) genes were analyzed using ReactomePA (39) and pathways were clustered into the major pathways from the Reactome database. Major pathways were filtered so that at least 12 sub-pathways were significantly enriched in at least one condition. (B) Dot plot showing top pathways enriched from Reactome database. Pathway coverage shown as gene ratio. –log_10_(p-value) presented as the dot size with WT STm in blue, T3SS-1^mut^ in green and T3SS-2^mut^ in red. Upregulated pathways shown on the right of the dotted line and down-regulated pathways on the left.

Few down-regulated pathways were observed at 2.5h pi, with more evident by 8h pi, largely related to cell cycle and DNA repair. Genes involved in cell cycle processes including checkpoints and mitotic (M) phase pathways were more highly suppressed in WT-infected HIOs, than in T3SS-1^mut^ and T3SS-2^mut^-infected HIOs **(Fig. 3B bottom)**. Taken together, our findings show that upregulated pathways primarily consisted of immune-related pathways that were only partially dependent on the two T3SS, while downregulated pathways dominated by cell cycle processes required both T3SS-1 and T3SS-2.

### Luminal STm contribute to rapid epithelial inflammatory gene expression

We also analyzed expression at the individual gene level, selecting pro-inflammatory gene sets from the Reactome database (cytokines, chemokines and antimicrobial peptides (AMPs)), to examine fold change relative to PBS-injected control HIOs **(Fig. 4A-C, Fig. S3)**. Induction of genes in all three categories occurred rapidly, characterized by markedly increased levels of cytokine, chemokine and AMP transcripts at 2.5h pi that were reduced by 8h pi. Global patterns revealed that infection with the T3SS-1^mut^ induced weaker stimulation of these proinflammatory mediators compared to the other infection conditions, although many transcripts were still up-regulated compared to PBS-injected HIOs. The strongest responders to infection were cytokines *CSF3*, also called granulocyte colony stimulating factor (G-CSF), and *IL17C*, and the antimicrobial peptide beta defensin-2 (*DEFB4*). Strong upregulation of *IL17C* and *DEFB4*, genes involved in epithelial intrinsic defenses (17–19), suggests that upon sensing infection, epithelial cells mount a direct antimicrobial response in addition to producing chemokines to recruit other immune cells. Notably, chemokine genes were not induced as strongly at these time points compared to cytokine and AMP genes **(Fig. 4B)**. Some other responses occurred independently of either T3SS-1 and T3SS-2 including Tumor Necrosis Factor (*TNF), IL8* and *CXCL5* as fold change was comparable between the three conditions while *IL6* expression was dependent on T3SS-1.

**Figure 4:**
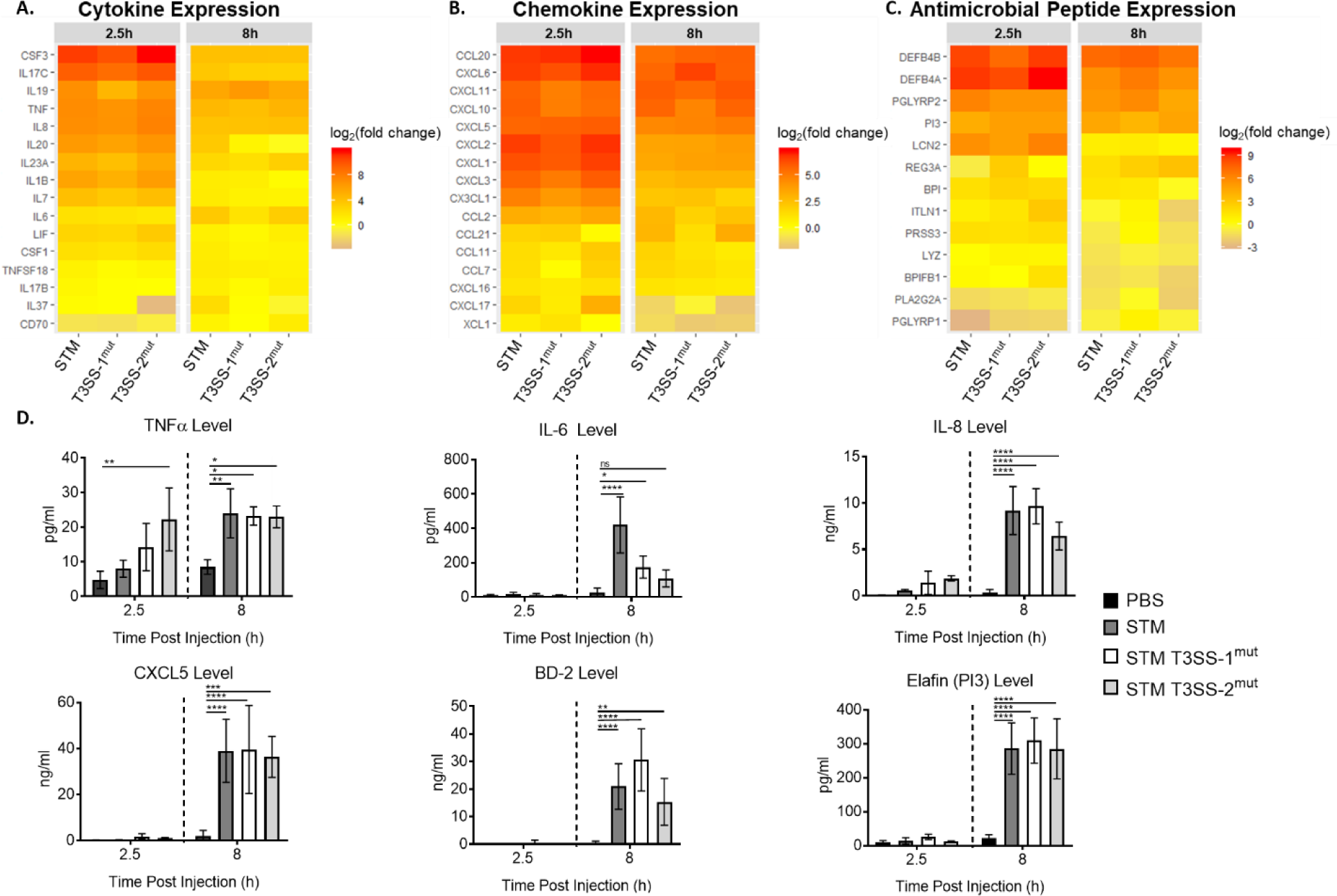
Cytokine, chemokine and antimicrobial peptide induction is not dependent on T3SS-1 or T3SS-2. (A-C) Gene expression presented as log_2_(fold change) relative to PBS injected HIOs at 2.5h and 8h post injection. (A) Cytokine expression, (B) Chemokine expression, (C) Antimicrobial peptide expression. (D) Cytokine, chemokine and antimicrobial peptide levels measured from HIO supernatant at 2.5 and 8h post injection via ELISA. n=4 biological replicates. Error bars represent SD. Significance calculated by two-way ANOVA.

To test whether gene level expression differences were reflected at the protein level, we collected supernatants from infected HIOs at 2.5h and 8h pi and measured cytokines by ELISA. In concordance with the transcript data, release of TNF, IL8, and CXCL5 were consistent across all three infection conditions **(Fig. 4D, Fig. S4)**. While the degree of transcript upregulation for AMPs varied between time points across the three infections, release of these mediators (Beta Defensin-2 and ELAFIN) into the medium did not significantly differ between WT and mutant infections. In contrast, IL6, which was increased in just STm and T3SS-2^mut^-infected HIOs by 8h pi at the transcriptional level, was present at significantly lower levels in supernatants from HIOs infected with either mutant. While there was less upregulation of *IL6* transcript in T3SS-1^mut^ infected HIOs compared to WT-infected HIOs, reduced levels of IL6 in the supernatant in T3SS-2^mut^ infected HIOs suggests there may be additional post-transcriptional regulation affecting IL6 production in the HIOs during T3SS-2^mut^ infection. Collectively, these results show that the HIOs mount a rapid pro-inflammatory, antimicrobial transcriptional response to STm infection and that invasion-defective T3SS-1^mut^ bacteria, previously reported to have a large defect in inducing an inflammatory response, can signal through the luminal compartment to induce robust inflammation following prolonged interactions with the epithelium.

### Downregulation of cell cycle pathways during STm infection is dependent on T3SS-1 and T3SS-2

Next, we turned our attention to genes that were downregulated during STm infection. Our pathway enrichment analysis identified cell cycle as the category containing the most significantly downregulated pathways. To further assess whether downregulation of cell cycle-related pathways was dependent on T3SS-1 and -2, we directly compared genes in the cell cycle pathway that were significantly changed in the three infection conditions. In agreement with our findings looking at major cellular processes responding to infection **(Fig. 3A)**, we found relatively few genes in the cell cycle pathway significantly downregulated compared to PBS-injected HIOs at 2.5h pi **(Fig. 5A)**. However, by 8h pi the number of significantly downregulated genes substantially increased from 76 genes to 161 genes in the WT-infected HIOs **(Fig. 5B)**. Gene downregulation was partially dependent on both T3SS-1 and -2, as only 68 genes and 58 genes, respectively, were significantly downregulated at 8h pi. These observations are consistent with a role for the T3SS in establishing an intracellular niche for STm replication.

**Figure 5:**
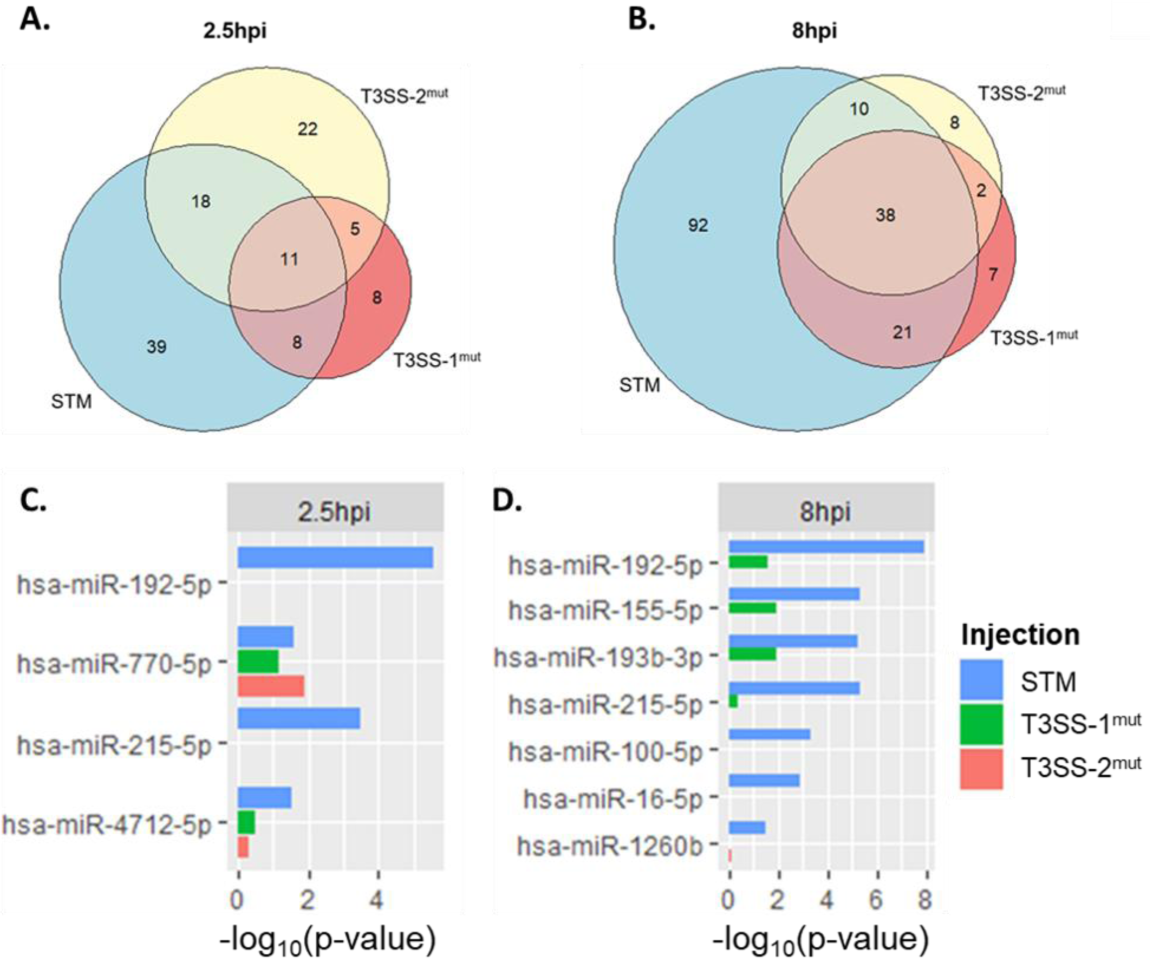
Cell cycle pathways are downregulated during STm infection dependent on T3SS-1 and T3SS-2. (A-B) Euler diagram comparison of cell cycle genes downregulated compared to PBS injected HIOs at 2.5h (A) and 8h (B) post injection. Genes were filtered by p-value < 0.05. (C-D) miRNA enrichment profiles were calculated using Gprofiler package in R (24) based on significantly downregulated genes compared to PBS injected HIOs. – log_10_(p-value) plotted for each miRNA that is significantly enriched in at least one infection condition.

Decreases in transcript levels can occur through several mechanisms, including halting synthesis of new transcripts or through degradation of existing transcripts by miRNA. Evidence for miRNA expression manipulation by pathogens, including *Salmonella*, continues to emerge (20–23), so we used gprofiler2 (24) as the basis for an informatics approach to identify potential regulatory miRNAs associated with our downregulated gene sets. Infection with WT STm resulted in the most significant predicted association of miRNAs, with elevated coverage of miRNA-regulated gene sets at 8h compared to 2.5h pi **(Fig. 5C-D)**. Notably, these miRNA species were not predicted to be strongly associated with the downregulated gene sets from T3SS-1^mut^ or T3SS-2^mut^-infected HIOs **(Fig. 5C-D)**. Several of these miRNA species including miR-192-5p and miR-155-5p that were more significantly associated with the WT-infected HIO gene set, are known to regulate cell proliferation (25, 26). These data suggest that miRNA-mediated downregulation of cell cycle genes may contribute to modulation of cell cycle-related pathways during *Salmonella* infection, in a T3SS-dependent manner.

## Discussion

Human intestinal epithelial responses to *Salmonella enterica* serovar Typhimurium are still incompletely understood, despite the prominent contribution of this species to human disease burden. Here we used the human intestinal organoid model to analyze transcriptional profiles defining early epithelial responses to STm infection, including the contribution of two major virulence determinants, T3SS-1 and -2. We found that HIOs responded rapidly and robustly to all 3 infections by upregulating pro-inflammatory pathways early and transiently, whereas downregulation of host pathways including cell cycle and DNA repair occurred later and only in WT STm-infected HIOs.

*Salmonella* infection strongly induces inflammatory responses, and exploits the inflammatory environment created during infection to outcompete the resident microbiota and replicate within the lumen of the intestine (27). Accordingly, our transcriptomics analysis found that the dominant response occurring in the HIOs was inflammatory. While this was largely expected for WT STm infection, based on studies in other model systems, we had predicted that infection with T3SS-1^mut^ would result in reduced activation of these pathways. Prior studies showed that T3SS-1 strongly contributes to the inflammatory response with significantly reduced levels of inflammation and colitis in mouse models, and little to no upregulation of pro-inflammatory cytokines in tissue culture models of STm infection (14–16). However, we observed largely similar patterns of induction of several pro-inflammatory mediators in HIOs infected with T3SS-1^mut^. This included IL8, which in HeLa cells was dependent on T3SS-1 effectors for upregulation (28). This finding highlights the advantage of using model systems that more closely reflect physiologic infection conditions. Although immortalized cell lines can more easily be manipulated than mouse models, the inoculum is removed after the initial infection, and therefore longer-term interactions between the luminal surface of the epithelium and the bacteria cannot be easily studied. The enclosed lumen of the HIOs naturally limits the extent of extracellular bacterial replication and allows study of these longer-term interactions, revealing a strong contribution of luminal bacteria in inducing upregulation of pro-inflammatory mediators since there was a >2-log defect in invasion with the T3SS-1^mut^.

Among our upregulated gene sets, key targets reflected known modulators of STm infection. The strongest responder in all 3 infection conditions, *CSF3* (encoding G-CSF), was previously implicated in regulating a super-shedder phenotype of *Salmonella* to enhance spread of the bacteria to other hosts, and injection of G-CSF in moderate-shedder animals recapitulated the super-shedder phenotype (29). Additionally, *IL17C* and *DEFB4* contribute to epithelial intrinsic defenses against bacterial pathogens through regulating intestinal barrier integrity and bacterial killing, respectively (17–19). Overall, the transcriptional responses across the 3 infection conditions were similar with only a slight decrease in upregulation in the T3SS-1^mut^-infected HIOs. Notably, upregulation of *IL6* expression appeared to be dependent on T3SS-1. Interestingly, while *IL6* transcript upregulation was dependent on T3SS-1, neither T3SS-1^mut^ or T3SS-2^mut^ infections stimulated significant IL6 protein production compared to PBS-injected HIOs. These observations suggest a novel function for T3SS-2 in post-transcriptional regulation of IL6 production. Together, these findings highlight several avenues for future study including IL6 post-transcription regulation by T3SS-2, and how *CSF3* regulation and function in the early stages of STm epithelial infection may contribute to a supershedder phenotype, using a HIO system reconstituted with immune cells, like neutrophils.

Down-regulation of host gene expression was dependent on T3SS-1 and -2, notably host cell cycle-related genes. Cell cycle regulation in the intestine affects the rate of cell turnover in order to shed infected or damaged cells and is therefore commonly targeted by bacteria (30, 31). A previous study from our consortium group showed that HIO colonization with a commensal strain of *E. coli* enhanced cell proliferation and could therefore be protective against invasive infections (6). In contrast, Holden and colleagues recently reported that STm can block cell cycle progression in mouse intestinal cells and proposed that this enhances intestinal colonization of STm (32). This study showed that T3SS-2 effectors regulated this phenotype through targeting proteins important for cleavage furrow formation, rather than exerting regulation at the transcriptional level. Here we found that that both T3SS-1 and T3SS-2 contribute to downregulating cell cycle-related transcripts, and this is the first study to our knowledge that implicates regulation of the cell cycle by STm at the transcript level. Moreover, miRNA expression is increasingly appreciated as a mechanism to regulate gene expression during bacterial infections and our results strongly predict regulation by specific miRNAs, opening an avenue for further exploration of cell cycle regulation by STm.

Collectively, the complex and dynamic transcriptional response in the STm-infected HIOs demonstrate the utility of using this non-transformed epithelial cell model to examine what aspects may be specific and physiologically relevant to human disease. HIOs supported both luminal and intracellular bacterial replication, while still maintaining overall structural integrity, better mimicking the interaction of both these bacterial populations with the epithelium *in vivo*. This model system allows for observation of infected cells, as well as bystander cells which can be studied using single cell RNA-seq, and because of the enclosed environment, the entire HIO can be visualized in sections or by live cell imaging. Additionally, with this enclosed lumen, it is possible to study sustained responses induced by the bacteria from the extracellular environment, an important aspect of STm infection biology that has been difficult to study in traditional tissue culture models. As further evidence to strengthen this model for future studies, our upregulated gene set for the WT infection was consistent with data from an earlier study, which looked at the HIO transcriptional response to WT STm infection (5), highlighting the reproducibility of this model system. Thus, this concordance opens areas for future work, including studying post-transcriptional regulation of cytokine production by T3SS-2 and transcriptional regulation of cell cycle processes by STm. Additionally, the HIO model is well suited to characterize host epithelial responses to other *Salmonella enterica* serovars in order to better understand how individual serovars interact uniquely with the host, as well as adding additional components such as a simplified microbiome or immune cells, to study more complex interactions between *Salmonella* and the human intestine.

## Materials and Methods

### HIO Differentiation and Culture

HIOs were generated by the In Vivo Animal and Human Studies Core at the University of Michigan Center for Gastrointestinal Research, as previously described (7,33). Human ES cell line WA09 (H9) was obtained from Wicell International Stem Cell Bank and cultured on Matrigel (BD Biosciences) coated 6-well plates in mTeSR1 media (Stem Cell Technologies) at 37°C in 5% CO2. Cells were seeded onto Matrigel-coated 24-well plates in fresh mTeSR1 media and grown until 85-90% confluence. Definitive endoderm differentiation was induced by washing the cells with PBS and culturing in endoderm differentiation media (RPMI 1640, 2%FBS, 2 mM L-glutamine, 100 ng/ml Activin A and 100 Units/ml Pen/Strep) for three days where media were exchanged each day. Cells were then washed with endoderm differentiation media without Activin A and cultured in mid/hindgut differentiation media (RPMI 1640, 2%FBS, 2 mM L-glutamine, 500 ng/ml FGF4, 500 ng/ml WNT3A and 100 Units/ml Pen/Strep) for 4 days until spheroids were present. Spheroids were collected, mixed with ice cold Matrigel (50μl of Matrigel + 25μl of media + 50 spheroids), placed in the center of each well of a 24-well plate, and incubated at 37°C for 10 minutes to allow Matrigel to solidify. Matrigel embedded spheroids were grown in ENR media (DMEM:F12, 1X B27 supplement, 2 mM L-glutamine, 100 ng/ml EGF, 100 ng/ml Noggin, 500 ng/mL Rspondin1, and 15 mM HEPES) for 14 days where medium was replaced every 4 days. Spheroids growing into organoids (HIOs) were dissociated from Matrigel by pipetting using a cut wide-tip (2-3 mm). HIOs were mixed with Matrigel (6 HIOs + 25μL of media + 50μL of Matrigel) and placed in the center of each well of 24-well plates and incubated at 37°C for 10 minutes. HIOs were further grown for 14 days in ENR media with medium exchanged every 4 days. Before use in experiments, HIOs were carved out of Matrigel, washed with DMEM:F12 media, and re-plated with 5 HIO/well in 50μL of Matrigel in ENR media with medium exchanged every 2-3 days for 7 days prior to microinjection.

### Bacterial Growth Condition and HIO Microinjection

STm strains used in this study are listed in Table S5. Strains were stored at −80°C in LB medium containing 20% glycerol and cultured on Luria-Bertani (LB, Fisher) agar plates. Selected colonies were grown overnight at 37°C under static conditions in LB liquid broth. Bacteria were pelleted, washed and re-suspended in PBS. Bacterial inoculum was estimated based on OD600 and verified by plating serial dilutions on agar plates to CFU. HIOs were cultured in groups of 5/well using 4-well plates (ThermoFisher). Individual HIO lumens were microinjected using a glass caliber needle with 1μl of PBS control or different STm mutants (10^5^CFU/HIO or 10^3^CFU/HIO for 24h infections). HIOs were washed with PBS and incubated for 2h at 37°C in ENR media to allow for re-sealing of the epithelial layer. HIOs were then treated with gentamicin (100 μg/ml) for 15 min to kill bacteria outside the HIOs, then incubated in fresh medium with gentamicin (10 μg/ml).

### Quantitative measurement of HIO-associated bacteria and cytokine secretion

Quantitation of viable bacteria was assessed per HIO. Individual HIOs were removed from Matrigel, washed with PBS and homogenized in PBS. Total CFU/HIO were enumerated by serial dilution and plating on LB agar. To assess intracellular bacterial burden, HIOs were sliced in half, treated with gentamicin (100 μg/ml) for 10 min to kill luminal bacteria, washed with PBS, homogenized and bacterial CFU were enumerated on LB-agar. Medium from each well (5 HIOs/well) was collected at indicated time points after microinjection and cytokines, chemokines and defensins were quantified by ELISA assay at the UM ELISA core.

### Immunohistochemistry and Immunofluorescence Staining

HIOs were fixed with 10% neutral buffered formalin or Carnoy’s solution for 2 days and embedded in paraffin. HIOs were sectioned (5 μm thickness) by the UM Histology Core and stained with hematoxylin and eosin (H&E). Carnoy’s-fixed HIO sections were stained with periodic acid-Schiff (PAS) staining reagents according to manufacturer’s instructions (Newcomersupply). H&E- and PAS-stained slides were imaged on an Olympus BX60 upright microscope. For immunofluorescence staining, formalin-fixed HIO sections were deparaffinized and subjected to antigen retrieval in sodium citrate buffer (10 mM Sodium citrate, 0.05% Tween 20, pH 6.0). Sections were permeabilized with PBS+ 0.2% Triton X-100 for 30 min, then incubated in blocking buffer (PBS, 5% BSA, and 10% normal goat serum) for 1h. Human Occludin was stained using rabbit anti-Occludin polyclonal antibody (ThermoFisher) in blocking buffer overnight at 4°C. Goat anti-mouse secondary antibody conjugated to Alexa-594 was used according to manufacturer’s instructions (ThermoFisher) for 1h RT in blocking buffer. *Salmonella* were stained using FITC-conjugated Anti-*Salmonella* Typhimurium antibody (Santa Cruz, 1E6) in blocking buffer for 1h RT. DAPI was used to stain DNA. Sections were mounted using coverslips (#1.5) and Prolong Diamond Antifade Mountant (ThermoFisher). Images were taken on the Nikon A1 confocal microscope and processed using ImageJ.

### RNA Sequencing and Analysis

Total RNA was isolated from groups of 5 HIOs per replicate with a total of 4 replicates per condition using the mirVana miRNA Isolation Kit (ThermoFisher). Cytosolic and mitochondrial ribosomal RNA was removed from samples using the Ribo-Zero Gold Kit according to manufacturer’s instructions (Illumina). The quality of RNA was confirmed (RIN >8.5) using a Bioanalyzer and used to prepare cDNA libraries by the UM DNA Sequencing Core. Libraries were sequenced on Illumina HiSeq 2500 platforms (single-end, 50 bp read length).

### Statistical Methods

Data were analyzed using Graphpad Prism 7 and R software. Differences between 2 groups were tested using the unpaired-t test or Mann-Whitney test. Differences between 3 or more groups were tested using Two-way ANOVA, followed by Tukey’s multiple comparisons test. The mean of at least 3 independent experiments was presented with error bars showing standard deviation (SD). P values of less than 0.05 were considered significant and designated by: *P < 0.05, **P < 0.01, ***P < 0.001 and ****P < 0.0001. All statistically significant comparisons within experimental groups are marked.

### Data and Software Availability

Data availability: deposition into ArrayExpress in progress. Source code for analyses can be found at: https://github.com/rberger997/HIO_dualseq2 and https://github.com/aelawren/HIO_RNAseq.

## RNAseq analysis protocol

### Sequence alignment

Sequencing generated FASTQ files of transcript reads were pseudoaligned to the human genome (GRCh38.p12) using kallisto software(34). Transcripts were converted to estimated gene counts using the tximport(35) package with gene annotation from Ensembl (36).

### Differential gene expression

Differential expression analysis was performed using the DESeq2 package (37) with p-values calculated by the Wald test and adjusted p-values calculated using the Benjamani & Hochberg method (38).

### Pathway enrichment analysis

Pathway analysis was performed using the Reactome pathway database and pathway enrichment analysis in R using ReactomePA software package (39). miRNA analysis was performed using Gprofiler2 package (24).

### Statistical analysis

Analysis was done using RStudio version 1.1.453. Plots were generated using ggplot2 (40) with data manipulation done using dplyr (41). Euler diagrams of gene changes were generated using the Eulerr package (42). Cluster heatmaps were generated using the pheatmap package (43).

## Acknowledgments

This work was supported by the NIAID U19AI116482-01 grant. A-LL. was supported by the Molecular Mechanisms of Microbial Pathogenesis training grant (NIH T32 AI007528). We thank the Host-Microbiome Initiative, the Center for Live Cell Imaging (CLCI), Microscopy and Image Analysis Laboratory (MIL), the Comprehensive Cancer Center Immunology and Histology Cores supported by the University of Michigan Cancer Center Support Grant (P30CA46592), and the DNA Sequencing Core at University of Michigan Medical School. We gratefully acknowledge members of the O’Riordan laboratory as well as members of the Wobus, Young, Takayama and Spence laboratories for many helpful discussions.

## References

1. 2020. Salmonella Homepage | CDC.

2. Valdez Y, Grassl GA, Guttman JA, Coburn B, Gros P, Vallance BA, Finlay BB. 2009. Nramp1 drives an accelerated inflammatory response during Salmonella-induced colitis in mice. Cell Microbiol 11:351–362.

3. Monack DM, Bouley DM, Falkow S. 2004. Salmonella typhimurium persists within macrophages in the mesenteric lymph nodes of chronically infected Nramp1+/+ mice and can be reactivated by IFNgamma neutralization. J Exp Med 199:231–241.

4. Spence JR, Mayhew CN, Rankin SA, Kuhar MF, Vallance JE, Tolle K, Hoskins EE, Kalinichenko VV, Wells SI, Zorn AM, Shroyer NF, Wells JM. 2011. Directed differentiation of human pluripotent stem cells into intestinal tissue in vitro. Nature 470:105–109.

5. Forbester JL, Goulding D, Vallier L, Hannan N, Hale C, Pickard D, Mukhopadhyay S, Dougan G. 2015. Interaction of Salmonella enterica serovar Typhimurium with intestinal organoids derived from human induced pluripotent stem cells. Infect Immun 83:2926–2934.

6. Hill DR, Huang S, Nagy MS, Yadagiri VK, Fields C, Mukherjee D, Bons B, Dedhia PH, Chin AM, Tsai Y-H, Thodla S, Schmidt TM, Walk S, Young VB, Spence JR. 2017. Bacterial colonization stimulates a complex physiological response in the immature human intestinal epithelium. Elife 6.

7. Leslie JL, Huang S, Opp JS, Nagy MS, Kobayashi M, Young VB, Spence JR. 2015. Persistence and toxin production by Clostridium difficile within human intestinal organoids result in disruption of epithelial paracellular barrier function. Infect Immun 83:138–145.

8. Jennings E, Thurston TLM, Holden DW. 2017. Salmonella SPI-2 Type III Secretion System Effectors: Molecular Mechanisms And Physiological Consequences. Cell Host Microbe 22:217–231.

9. Lou L, Zhang P, Piao R, Wang Y. 2019. Salmonella Pathogenicity Island 1 (SPI-1) and Its Complex Regulatory Network. Front Cell Infect Microbiol 9:270.

10. Alteri CJ, Himpsl SD, Pickens SR, Lindner JR, Zora JS, Miller JE, Arno PD, Straight SW, Mobley HLT. 2013. Multicellular Bacteria Deploy the Type VI Secretion System to Preemptively Strike Neighboring Cells. PLoS Pathogens.

11. Galan JE, Curtiss RY Iii. allow Salmonella typhimurium to penetrate tissue culture cells.

12. Pfeifer CG, Marcus SL, Steele-Mortimer O, Knodler LA, Finlay BB. 1999. Salmonella typhimurium virulence genes are induced upon bacterial invasion into phagocytic and nonphagocytic cells. Infect Immun 67:5690–5698.

13. Laughlin RC, Knodler LA, Barhoumi R, Payne HR, Wu J, Gomez G, Pugh R, Lawhon SD, Bäumler AJ, Steele-Mortimer O, Adams LG. 2014. Spatial segregation of virulence gene expression during acute enteric infection with Salmonella enterica serovar Typhimurium. MBio 5:e00946–13.

14. Bruno VM, Hannemann S, Lara-Tejero M, Flavell RA, Kleinstein SH, Galán JE. 2009. Salmonella Typhimurium type III secretion effectors stimulate innate immune responses in cultured epithelial cells. PLoS Pathog 5:e1000538.

15. Hapfelmeier S, Stecher B, Barthel M, Kremer M, Müller AJ, Heikenwalder M, Stallmach T, Hensel M, Pfeffer K, Akira S, Hardt W-D. 2005. The Salmonella pathogenicity island (SPI)-2 and SPI-1 type III secretion systems allow Salmonella serovar typhimurium to trigger colitis via MyD88-dependent and MyD88-independent mechanisms. J Immunol 174:1675–1685.

16. Barthel M, Hapfelmeier S, Quintanilla-Martínez L, Kremer M, Rohde M, Hogardt M, Pfeffer K, Rüssmann H, Hardt W-D. 2003. Pretreatment of mice with streptomycin provides a Salmonella enterica serovar Typhimurium colitis model that allows analysis of both pathogen and host. Infect Immun 71:2839–2858.

17. Cobo ER, Chadee K. 2013. Antimicrobial Human β-Defensins in the Colon and Their Role in Infectious and Non-Infectious Diseases. Pathogens 2:177–192.

18. Reynolds JM, Martinez GJ, Nallaparaju KC, Chang SH, Wang Y-H, Dong C. 2012. Cutting edge: regulation of intestinal inflammation and barrier function by IL-17C. J Immunol 189:4226–4230.

19. Gong H, Ma S, Liu S, Liu Y, Jin Z, Zhu Y, Song Y, Lei L, Hu B, Mei Y, Liu H, Liu Y, Wu Y, Dong C, Xu Y, Wu D, Liu H. 2018. IL-17C Mitigates Murine Acute Graft-vs.-Host Disease by Promoting Intestinal Barrier Functions and Treg Differentiation. Front Immunol 9:2724.

20. Aguilar C, Cruz AR, Rodrigues Lopes I, Maudet C, Sunkavalli U, Silva RJ, Sharan M, Lisowski C, Zaldívar-López S, Garrido JJ, Giacca M, Mano M, Eulalio A. 2020. Functional screenings reveal different requirements for host microRNAs in Salmonella and Shigella infection. Nat Microbiol 5:192–205.

21. Herrera-Uribe J, Zaldívar-López S, Aguilar C, Luque C, Bautista R, Carvajal A, Claros MG, Garrido JJ. 2018. Regulatory role of microRNA in mesenteric lymph nodes after Salmonella Typhimurium infection. Vet Res 49:9.

22. Schulte LN, Eulalio A, Mollenkopf H-J, Reinhardt R, Vogel J. 2011. Analysis of the host microRNA response to Salmonella uncovers the control of major cytokines by the let-7 family. EMBO J 30:1977–1989.

23. Huang T, Huang X, Chen W, Yin J, Shi B, Wang F, Feng W, Yao M. 2019. MicroRNA responses associated with Salmonella enterica serovar typhimurium challenge in peripheral blood: effects of miR-146a and IFN-γ in regulation of fecal bacteria shedding counts in pig. BMC Vet Res 15:195.

24. Raudvere U, Kolberg L, Kuzmin I, Arak T, Adler P, Peterson H, Vilo J. 2019. g:Profiler: a web server for functional enrichment analysis and conversions of gene lists (2019 update). Nucleic Acids Res 47:W191–W198.

25. Yan-Chun L, Hong-Mei Y, Zhi-Hong C, Qing H, Yan-Hong Z, Ji-Fang W. 2017. MicroRNA-192-5p Promote the Proliferation and Metastasis of Hepatocellular Carcinoma Cell by Targeting SEMA3A. Appl Immunohistochem Mol Morphol 25:251–260.

26. Qu Y-L, Wang H-F, Sun Z-Q, Tang Y, Han X-N, Yu X-B, Liu K. 2015. Up-regulated miR-155-5p promotes cell proliferation, invasion and metastasis in colorectal carcinoma. Int J Clin Exp Pathol 8:6988–6994.

27. Winter SE, Thiennimitr P, Winter MG, Butler BP, Huseby DL, Crawford RW, Russell JM, Bevins CL, Adams LG, Tsolis RM, Roth JR, Bäumler AJ. 2010. Gut inflammation provides a respiratory electron acceptor for Salmonella. Nature 467:426–429.

28. Figueiredo JF, Barhoumi R, Raffatellu M, Lawhon SD, Rousseau B, Burghardt RC, Tsolis RM, Bäumler AJ, Adams LG. 2009. Salmonella enterica serovar Typhimurium-induced internalization and IL-8 expression in HeLa cells does not have a direct relationship with intracellular Ca(2+) levels. Microbes Infect 11:850–858.

29. Gopinath S, Hotson A, Johns J, Nolan G, Monack D. 2013. The systemic immune state of super-shedder mice is characterized by a unique neutrophil-dependent blunting of TH1 responses. PLoS Pathog 9:e1003408.

30. MacDonald TT. 1992. Epithelial proliferation in response to gastrointestinal inflammation. Ann N Y Acad Sci 664:202–209.

31. El-Aouar Filho RA, Nicolas A, De Paula Castro TL, Deplanche M, De Carvalho Azevedo VA, Goossens PL, Taieb F, Lina G, Le Loir Y, Berkova N. 2017. Heterogeneous Family of Cyclomodulins: Smart Weapons That Allow Bacteria to Hijack the Eukaryotic Cell Cycle and Promote Infections. Front Cell Infect Microbiol 7:208.

32. Santos AJM, Durkin CH, Helaine S, Boucrot E, Holden DW. 2016. Clustered Intracellular Salmonella enterica Serovar Typhimurium Blocks Host Cell Cytokinesis. Infect Immun 84:2149–2158.

33. McCracken KW, Howell JC, Wells JM, Spence JR. 2011. Generating human intestinal tissue from pluripotent stem cells in vitro. Nat Protoc 6:1920–1928.

34. Bray NL, Pimentel H, Melsted P, Pachter L. 2016. Near-optimal probabilistic RNA-seq quantification. Nat Biotechnol 34:525–527.

35. Soneson C, Love MI, Robinson MD. 2015. Differential analyses for RNA-seq: transcript-level estimates improve gene-level inferences. F1000Res 4:1521.

36. Rainer J. 2017. EnsDb.Hsapiens.v75: Ensembl based annotation package. R package version 2.99.0.

37. Love MI, Huber W, Anders S. 2014. Moderated estimation of fold change and dispersion for RNA-seq data with DESeq2. Genome Biol 15:550.

38. Benjamini Y, Hochberg Y. 1995. Controlling the False Discovery Rate: A Practical and Powerful Approach to Multiple Testing. Journal of the Royal Statistical Society: Series B (Methodological).

39. Yu G, He Q-Y. 2016. ReactomePA: an R/Bioconductor package for reactome pathway analysis and visualization. Mol Biosyst 12:477–479.

40. Wickham H. 2016. ggplot2: Elegant Graphics for Data Analysis. Springer.

41. Wickham H, Francois R, Henry L, Müller K, Others. 2015. dplyr: A grammar of data manipulation. R package version 0 4 3.

42. Larsson J. 2019. eulerr: Area-Proportional Euler and Venn Diagrams with Ellipses.

43. Kolde R. 2018. pheatmap: Pretty Heatmaps.

44. Jones BD, Falkow S. 1994. Identification and characterization of a Salmonella typhimurium oxygen-regulated gene required for bacterial internalization. Infect Immun 62:3745–3752.

45. Guy RL, Gonias LA, Stein MA. 2000. Aggregation of host endosomes by Salmonella requires SPI2 translocation of SseFG and involves SpvR and the fms– aroE intragenic region. Mol Microbiol.

